# Improving conformational ensembles of folded proteins in GōMartini

**DOI:** 10.1101/2025.10.24.684331

**Authors:** Maksim Kalutskii, Carter J. Wilson, Helmut Grubmüller, Maxim Igaev

## Abstract

The Martini coarse-grained (CG) force field enables efficient simulations of biomolecular systems but cannot reliably maintain folded protein structures. To stabilize proteins during simulation, Martini is typically combined with structure-based force fields such as elastic network models (ENMs) or Gō models. While these approaches preserve global folds and capture protein flexibility, their ability to reproduce conformational dynamics remains unclear. Here, we benchmark Martini combined with ENMs or Gō models on three folded proteins and show that both approaches struggle to sample the conformational space observed in atomistic simulations, even when uniform interaction strengths or equilibrium bond distances are adjusted. This limitation arises from the assumption of a uniform interaction network, in which all bond energies are equal. To overcome this, we present a fully automated, perturbation-based optimization approach for Gō networks, *PoGō*, that iteratively refines a non-uniform Gō network against a pre-computed atomistic free energy landscape in essential conformational space. Our approach converges rapidly, yielding CG ensembles in close agreement with reference atomistic simulations. As a cross-validation, the optimization also improves the root-mean-square fluctuation profile.

## Introduction

Proteins perform a vast array of fundamental functions in living systems, including catalyzing reactions, ^1,2^ facilitating protein folding, ^3,4^ signal transduction, ^5,6^ and transporting molecules. ^7,8^ These processes often require proteins to undergo conformational changes, allowing them to alternate between biologically functional states. ^9,10^ As more protein structures are deposited into the Protein Data Bank and more molecular simulations are performed, a key observation has emerged: the majority of protein dynamics can be described by a surprisingly small number of collective variables which can be effectively identified using dimensionality reduction techniques such as principal component analysis (PCA), ^12,13^ full correlation analysis (FCA), ^14^ time-independent component analysis (TICA), ^15^ functional mode analysis (FMA), ^16,17^ or coevolution-driven analysis. ^18^ A recent review ^19^ provides a more comprehensive summary of conventional and state-of-the-art machine learning methods for inferring collective variables from molecular simulations.

When applied to all-atom molecular dynamics (MD) trajectories, PCA usually yields a small fraction of principal components (PCs) that capture most of the ensemble variance. ^12,13^ Projecting the high-dimensional dynamics along these “essential” PCs and quantifying the low-dimensional free energy landscape is key to our understanding of protein conformational dynamics and folding. ^12,20–26^ Enhanced sampling techniques can further exploit this low-dimensional picture of protein dynamics to accelerate exploration of the free energy landscape. ^27–31^ This low-dimensional framework also suggests that the details of fast degrees of freedom may be less essential for characterizing critical protein functions over extended timescales. This insight has inspired the development of coarse-grained (CG) models of proteins and complexes thereof, ^32–36^ which abandons the explicit description of fast degrees of freedom in favor of a simplified yet sufficiently accurate representation of the system of interest. In some cases, the essential dynamics of a protein have even been used to directly guide CG topology design. ^37^

Coarse-graining presents a compelling approximation to atomistic MD because it significantly reduces computational costs, thus enabling improved sampling and access to spatiotemporal scales that are often computationally infeasible with all-atom approaches. The Martini force field ^32,38^ is one of the most widely used physics-based CG force fields for simulating biomolecular systems. However, due to the averaging of directional interactions, such as hydrogen bonds, Martini is unable to maintain folded protein structure; in light of this limitation, structure-based CG models, such as elastic network models (ENMs) ^39^ or Gō models, ^11,40^ are often used in combination with Martini. ^41–43^ While ENMs can accurately capture local protein flexibility, they are unable to describe large conformational changes due to the harmonic nature of their bonds. ^41^ To overcome this limitation and thereby to also increase protein flexibility, including partial unfolding, Gō models with breakable bonds encoded as Lennard-Jones potentials are usually employed instead ^41–44^

One of the main shortcomings of using structure-based force fields for generating protein conformational ensembles is that these are typically unable to correctly describe the correct distribution of variance across PCs. ^45^ Whereas atomistic MD simulations combined with PCA typically identify a few soft, highly collective motions that capture most of the variance, ^12,13^ both ENMs and Gō models tend to spread the same variance across a much larger number of PCs. ^45,46^ As a result, the free energy landscape in the essential subspace can deviate considerably from that produced by atomistic force fields. Although combining Martini with the Gō model seems to be a more promising approach for studying large conformational changes in CG simulations, ^43,47^ it still samples only a small fraction of atomistic conformations, even when optimized to reproduce atomistic fluctuations. ^48^

Recently, it has been proposed to decouple the development of Martini from that of the Gō model and the ENM. ^43^ Specifically, the Martini development should follow a standard building block optimization strategy, while the Gō model or the ENM should only be employed to bias the simulated ensemble based on experimental or all-atom MD data. ^43^ Along these lines, we suggest that fine-tuning structure-based potentials should seek to improve the agreement between atomistic and CG ensembles, with the goal of reproducing the free energy landscape along the essential PCs, rather than only matching fluctuation profiles, that neglect higher correlations, or identifying heuristic hyper-parameters that are fit using training datasets. ^43^

Optimizing the free-energy landscape in the essential subspace is a particularly challenging task. Standard methods for optimizing coarse-grained force fields, such as iterative Boltzmann inversion ^49^ or relative entropy minimization, ^50^ require re-running sufficiently long CG simulations at every optimization step to resample the free-energy landscape, which makes these approaches computationally intractable for high-dimensional parameter spaces.

In this paper, we develop and assess *PoGō*, a perturbation-based optimization method for Gō networks, here specifically applied to GōMartini. This framework aims to estimate the effect of network improvements on the conformational ensemble without performing explicit MD simulations, hence facilitating fast optimization. When combined with the particle swarm optimization (PSO) algorithm, ^51,52^ our approach yields force field perturbations that drastically improve the agreement between a CG and an atomistic reference ensemble.

We tested our method on three different protein systems, and obtained converged Gō networks within tens of optimization steps in each case. The resulting CG free energy landscapes in the essential subspace agree well with those of the corresponding atomistic reference simulations. Moreover, we find that, while not explicitly optimizing atomistic fluctuations, improving agreement along the essential PCs also improves the agreement between the atomistic and CG fluctuation profiles. In short, we provide a fully automated approach for optimizing the essential dynamics and fluctuations of a GōMartini-based protein model.

## Methods

### Perturbation theory

Following Koyama *et al*., ^53^ we formulate the expression for the change in the conformational distribution of a simulated protein induced by an arbitrary set of linearly independent perturbation functions.

Consider a molecular system described by the additive potential function 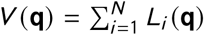, where **q** are the Cartesian coordinates. Measuring all energies in units of *k*_*B*_*T*, where *k*_*B*_ is the Boltzmann constant and *T* is the temperature, its canonical distribution is written as

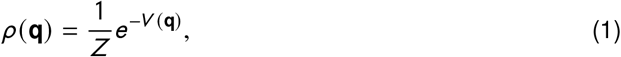

where *Z* = ∫*e*^−*V*^ *d* ^3*N*^ *q* is the partition function.

From the set of *N* potentials constituting *V* (**q**), we choose a subset of *M* ≤ *N* potentials **L**(**q**) = [*L*_1_ (q), *L*_2_ (**q**), …, *L*_*M*_ (**q**)] ^*T*^ and define perturbation coefficients ***λ*** = [*λ*_1_, *λ*_2_, …, *λ* _*M*_]^*T*^ to construct a new, perturbed potential

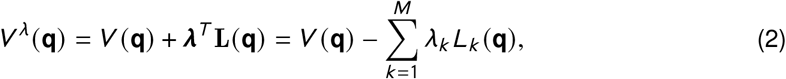

such that the canonical distribution corresponding to this perturbed potential is given by

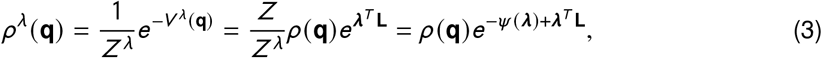

where we define 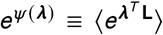, and ⟨… ⟩ denotes the statistical average with respect to the unperturbed distribution *ρ* (**q**).

With this perturbation, the interaction corresponding to one of the selected potentials *L*_*k*_ (**q**), where *k* ∈ {1, …, *M* }, is strengthened when *λ*_*k*_ > 0 and it is weakened when *λ*_*k*_ < 0. When ***λ* = 0**, the original and perturbed potentials are identical.

Since we are only interested in small perturbations, *i*.*e*. ‖***λ*** ‖ ≡ ***λ***^*T*^ ***λ*** ≪ 1, we use a second-order Taylor expansion of *ψ* (***λ***) at ***λ* = 0** yielding

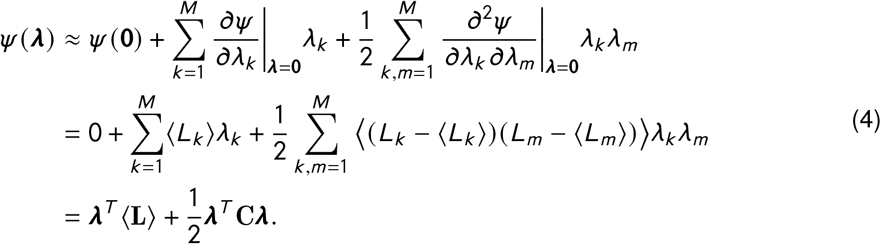

By using this approximate expression, Eq. 3 reads

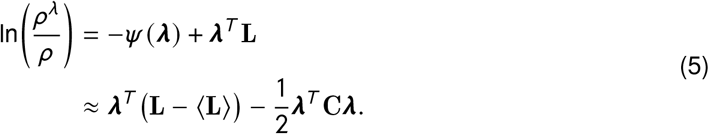

We note that the covariance matrix **C** is positive semi-definite and, therefore, can be represented as **C = UΩU**^*T*^, where **U** = [**u**_1_, …, **u**_*M*_] are the orthonormal eigenvectors of **C** and 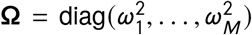 is the diagonal matrix that contains the corresponding non-negative eigenvalues. By performing the change of basis ***λ* = U*ξ*** and defining **B ≡ U**^*T*^ (**L −** ⟨**L**⟩), the following approximate expression for the change in the conformational distribution induced by the perturbation in Eq. 2 is obtained,

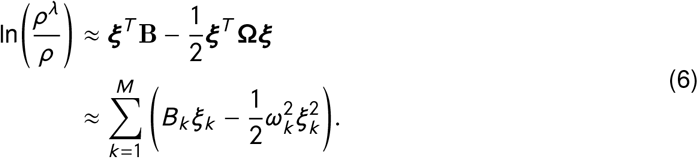

Equation 6 provides a computationally efficient way to quantify how small perturbations to linear interaction terms affect an equilibrium conformational distribution.

### Ensemble similarity metrics

In what follows we introduce three metrics to quantify the agreement between the essential subspaces of the tested atomistic and CG models.

The *root mean square inner product*

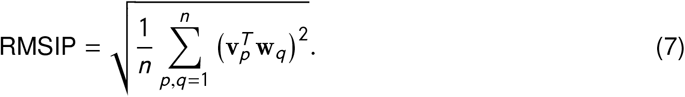

measures the geometric similarity between two subspaces spanned by the first *n* eigenvectors, {**v**_1_, … **v**_*n*_ } and {**w**_1_ … **w**_*n*_ }, corresponding to the two ensembles:

The *covariance overlap*

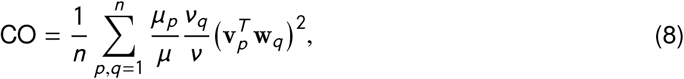

is similar to RMSIP but emphasizes subspace overlaps along directions of large variance, while downweighting overlaps that carry little variance. Here, 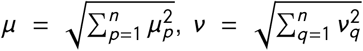 and {*µ*_1_, … *µ*_*n*_ } and {*ν*_1_, …, *ν*_*n*_ } are the respective eigenvalues. Note that both RMSIP and CO range from 0 (no overlap) to 1 (perfect agreement).

The *sliced Wasserstein distance* (SWD) of order *γ* ∈ [1, ∞),

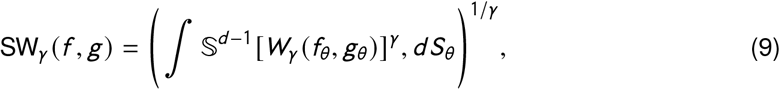

is a computationally efficient approximation of the full Wasserstein distance

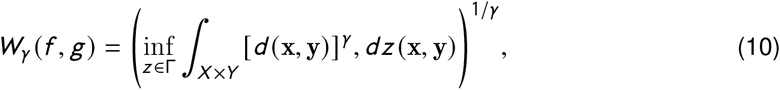

of the same order between two probability distributions, *f* on *X* ⊂ ℝ^*d*^ and *g* on *Y* ⊂ ℝ^*d*^. Here, *d* (#x00B7;, #x00B7;) is the distance metric on *X* × *Y* (i.e., the transportation cost), and Γ denotes the set of all possible joint distributions *z* (transportation plans) whose marginals are *f* and *g*. ^54^ Because computing the WD directly becomes prohibitively expensive for high-dimensional spaces (*d* > 1), the SWD offers a tractable alternative. It approximates the WD by projecting both distributions onto one-dimensional subspaces (or “slices”) defined by directions *θ* sampled uniformly from the unit sphere *S*^*d* −1^. *W*_*γ*_ (*f*_*θ*_, *g*_*θ*_) is the one-dimensional Wasserstein distance between the projected distributions *f*_*θ*_ and *g*_*θ*_, and *d S*_*θ*_ is the uniform measure on *S*^*d* −1^.

Unlike other structural similarity metrics such as RMSIP or CO, the SWD provides a true distance measure that ranges from 0 (perfect agreement) to +∞ (no overlap). Unless otherwise stated, we use SWD of order *γ* = 3 in all analyses.

### All-atom MD and Martini CG simulations

All molecular dynamics simulations were performed using GROMACS 2023. ^55^ For the atomistic simulations, the Amber99SB^*^-ILDN ^56^ force field and TIP3P water model ^57^ was used; for the CG simulations, Martini 3 was used. ^32^ Martini topology generation was performed using (`martinize2`. ^58^ Initial elastic (*κ* = = 500 kJ/mol/nm) and Gō networks (ϵ = 9.414 kJ/mol) were constructed using reference crystal structures, i.e., T4 lysozyme: PDB ID 182L, E. coli ribose binding protein: PDB ID 2DRI, and E. coli maltose binding protein: PDB ID 1MPB, with default cutoffs i.e., ENM: [0, 0.9 nm] and Gō: [0.3, 1.1 nm]. These PDB structures were used for both coarse-grained and atomistic simulations. For all systems, an initial minimization was performed using the steepest descent algorithm. For all production simulations the leap-frog integrator was used with a timestep of 2 fs for the atomistic and 20 fs for the CG systems. A temperature of 300 K was maintained using the velocity-rescaling thermostat ^59^ with a 1 ps coupling time and the pressure was maintained at 1 bar using the C-rescale barostat ^60^ with a coupling time of 5 ps. For the atomistic simulations, long-range electrostatic interactions were calculated using the Particle-mesh Ewald method ^61^ with a real-space cut-off of 1.0 nm and grid spacing of 0.12 nm, while the Lennard-Jones interactions were truncated at 1.0 nm and a dispersion correction was applied. Bonds to hydrogen atoms were constrained using the Parallel LINear Constraint Solver. ^62^ For the CG simulations, electrostatics were treated using a reaction-field with a 1.1 nm cutoff and relative dielectric of *ϵ* = 15. Lennard-Jones interactions were modified with a potential-shift and a cutoff of 1.1 nm.

## Results and discussion

### Limitations of uniform Gō networks in reproducing essential dynamics

To assess the accuracy of the essential dynamics relative to all-atom simulations we chose three test systems: T4 lysozyme (T4L), *E. coli* ribose binding protein (RBP), and *E. coli* maltose binding protein (MBP). We produced 5 × 600 ns of all-atom MD trajectories as well as Martini sampling using an ENM or a Gō network. We discarded the first 100 ns from each trajectory as an equilibration phase (see Methods). The atomistic trajectories were forward-mapped into a Martini representation using `martinize2`. ^58^

We then performed Cartesian-space PCA on the backbone beads of the forward-mapped atomistic trajectories for each of the three test systems. Projecting the same atomistic trajectories along the first three modes revealed diverse characteristic motions, consistent with previous reports. Specifically, the analysis of the T4L trajectories identified a hinge-bending motion (PC1), a twisting motion (PC2), and a torsional domain motion (PC3), which accounted for 21%, 15%, and 8% of the total ensemble variance, repsectively (Fig. 1a,b). Similarly, RBP exhibited a domain-closing motion (PC1), a twisting motion (PC2), and a propeller motion (PC3), capturing 31%, 30%, and 11% of the total ensemble variance, respectively. For MBP, a domain-closing motion (PC1), a twisting motion (PC2), and a helix shift (PC3) explained 20%, 15%, and 9% of the variance, respectively (Fig. S1).

**Figure 1:**
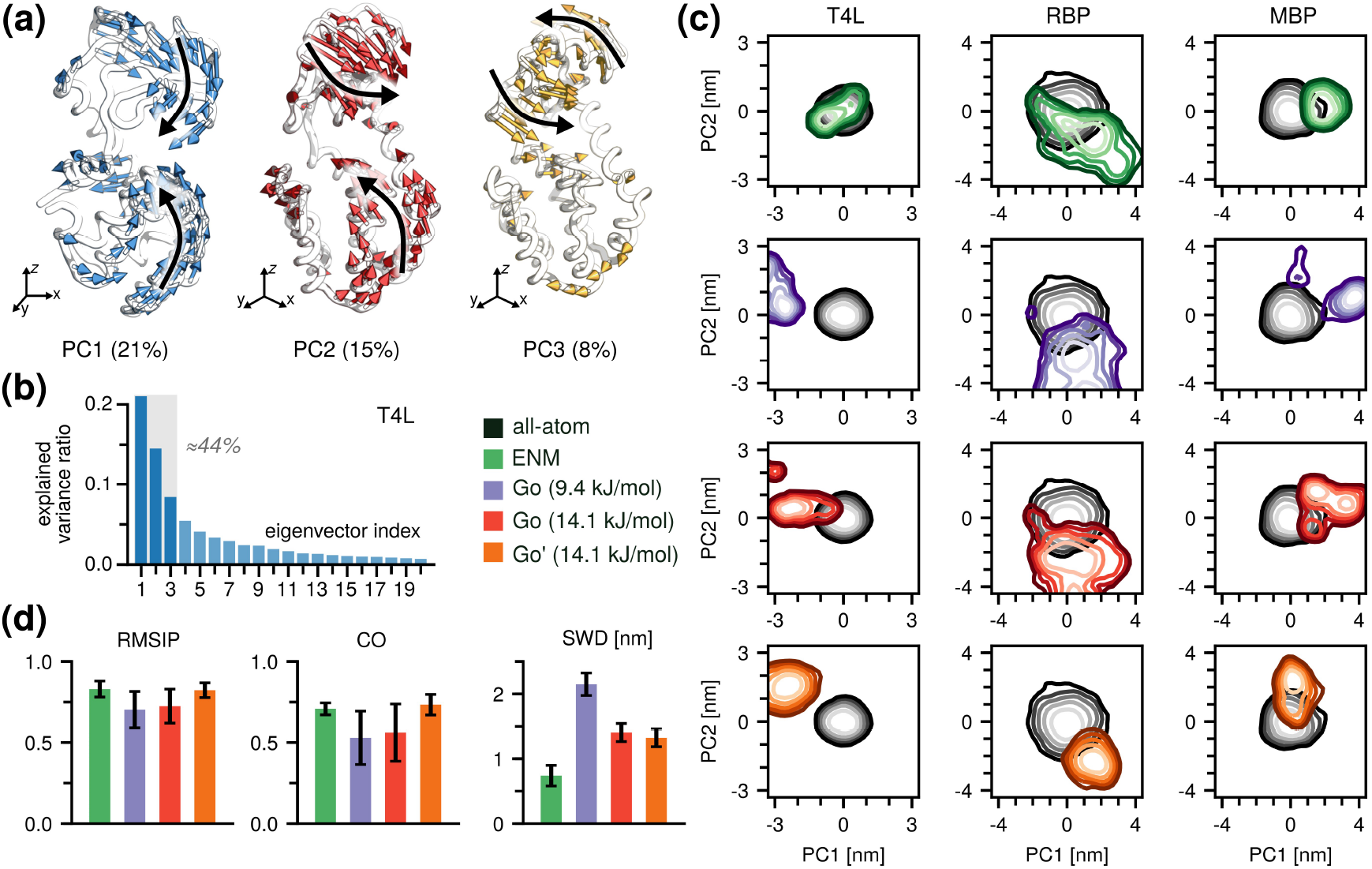
Performance of unoptimized GōMartini. **(a)** The first three PC modes of T4L and their contributions to the total variance. **(b)** The first 20 eigenvalues sorted in descending order. **(c)** All-atom MD and CG trajectories projected onto the essential subspace of the MD ensembles (PC1 and PC2). Each column corresponds to a different protein, and each row corresponds to a different CG setup. **(d)** Root mean squared inner product (RMSIP), covariance overlap (CO) and sliced Wasserstein distance (SWD) calculated between the all-atom MD and CG ensembles in the essential subspace of the MD ensembles.

Projections of both the all-atom and CG trajectories onto the essential subspace revealed that the uniform, 9.4 kJ/mol Gō model ensembles were markedly more expanded than the atomistic ones and, in the case of T4L and MBP, sampled multiple minima (Fig. 1c and Fig. S2). The ENM ensembles of T4L and RBP exhibited distorted but partially overlapped with the atomistic distributions, while MBP showed a much tighter distribution (Fig. 1c and Fig. S2). Overall, neither the ENM nor the uniform 9.4 kJ/mol Gō model fully reproduced the sampling of the all-atom MD simulations within the essential subspace.

To compare the essential dynamics captured by the all-atom and CG models more systematically, we employed three complementary metrics: the root mean square inner product (RMSIP) (Eq. 7), the covariance overlap (CO) (Eq. 8), and the sliced Wasserstein distance (SWD) (Eq. 9, see Methods). For clarity, we only report the means and standard errors for each metric calculated over all simulation replicas and all protein systems.

The average RMSIP was 0.83 ± 0.05 for the ENM and 0.70 ± 0.11 for the Gō model, while we calculated the average CO of 0.71 ± 0.03 for the ENM and 0.53 ± 0.16 for the Gō model (Fig. 1d, left and middle). This result indicates that both models are able to capture reasonably well the overall essential modes of the test proteins and overall shape of the essential subspace.

Although both models reproduce the overall shape and dominant directions of the essential subspace, our primary interest lies in how the explored conformational space is populated. RMSIP and CO primarily measure the geometric similarity between distinct subspaces, focusing on whether the dominant directions of motion align, but they are largely insensitive to how the system samples along those directions. In particular, they do not account for differences in the mean position of the sampled distributions and consider only the relative magnitudes of the eigenvalues. As a result, they may yield high similarity scores even when the underlying populations of conformational states show little or no overlap. In this respect, the SWD provides a better metric to compare the sampled distributions themselves. The calculated SWD values confirmed these observations, yielding 0.83 ± 0.18,nm for the ENM and a much larger distance of 2.07 ± 0.16,nm for the Gō model, mainly due to the poor overlap for T4L (Fig. 1d, right side).

For all three test proteins, but particularly for T4L, we observed that the ENM outperformed the standard Gō model with respect to reproducing the atomistic sampling within the essential subspace. We attribute the better performance of the ENM to the harmonic nature of its bonds. In particular, harmonic and unbreakable bonds of the ENM likely restrict significant deviations from the initial conformation, while the breakable Gō bonds initialized with a relatively weak *ε* likely cause distortions of the reference structure.

To assess whether strengthening the Gō network could improve its performance, we increased the interaction strength by 50% (*ε* = 14.1 kJ/mol). This adjustment led to a contraction of the sampled ensemble and an improved SWD which shifted from 2.07 ± 0.16 nm to 1.38 ± 0.13 nm. We observed an small improvement in RMSIP and no significant change in CO.

We next investigated the effect of modifying the equilibrium Lennard-Jones distances of the Gō potentials, *i*.*e*., *r*_min,*i*_ = 2^1/6^*σ*_*i*_. Specifically, we calculated the average distance for each pair of backbone beads *i* and *j* (*r*_*ij*_) from the atomistic trajectory and set it as an equilibrium distance for the corresponding Gō potential, here denoted Gø’. This adjustment further improved the CO from 0.56 ± 0.18 to 0.73 ± 0.06 and produced smaller improvements in RMSIP and SWD.

These results suggest that (i) Martini simulations with default ENM or Gō networks do not accurately reproduce the free energy landscape in the essential space observed by all-atom MD, and that (ii) uniformly increasing the strength of the Gō potentials or adjusting their equilibrium distances results in a modest yet suboptimal improvement.

### Perturbation-based optimization of non-uniform Gō networks

These results suggest(Fig. 2a) that optimizing a non-uniform network of Gō potentials is required to match a reference ensemble in the essential subspace. Here, we outline a method for optimizing a non-uniform network using the perturbation theory framework described in the Methods. The atomistic ensemble is first mapped into the Martini CG representation, after which a principal component analysis (PCA) is performed and the trajectory is projected onto the resulting essential subspace to obtain the target distribution, *ρ*_AA_(**q**). A CG ensemble is then generated, initialized with a uniform Gō network, and then projected onto the same subspace to obtain *ρ*_CG_(**q**), which serves as the basis for optimization (Step 1 in Fig. 2a).

**Figure 2:**
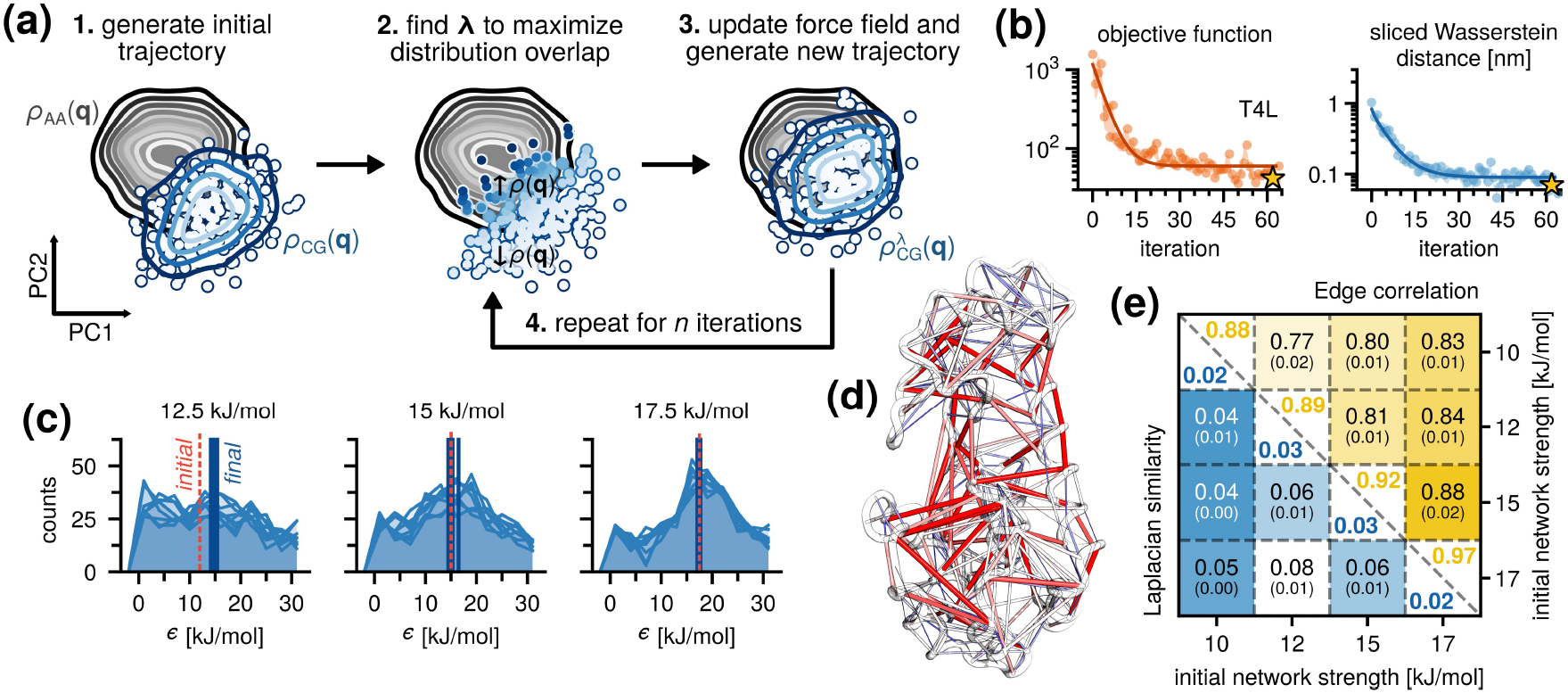
Perturbation-based optimization algorithm. **(a)** Sampling overlap between a CG ensemble (blue) and the reference all-atom MD ensemble (black) in the essential subspace. Given an initial CG ensemble and probability distribution (contour lines) (Step 1), the goal is to find a perturbation ***λ*** that increases the probability of overlapping configurations, ↑ *ρ* (**q**), and decreases the probability of non-overlapping configurations, ↓ *ρ* (**q**) (Step 2). With the optimal ***λ***, a new trajectory is generated based on the modified potential (Step 3), followed by finding the next optimal perturbation (Step 4). **(b)** Convergence behavior of the objective function and SWD for T4L. The iteration with the best agreement is indicated with a star. **(c)** Distribution of the optimized Gō network strengths for multiple replicas and for different initial (uniform) strengths. The initial and average values are indicated with vertical lines. **(d)** A representative optimized network mapped onto the 3D structure of T4L in the CG representation. Edge colors and sizes reflect the strength of the interaction. **(e)** Comparison of the optimized Gō networks for different initial (uniform) strengths. Values along the diagonal correspond to the networks whose optimizations started with the same initial strength. Standard errors are provided in brackets.

We then construct the following objective function to be minimized:

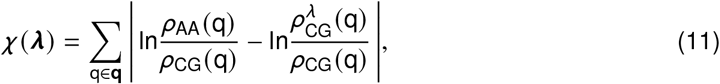

where 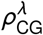 (**q**) is the projection of the canonical distribution for the perturbed CG ensemble (see Methods, Eq. (3)). The advantage of this objective function is that the expression ln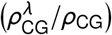 can be computed using the approximate analytical expression in Eq. 5 that only depends on ***λ*** and *ρ*_CG_(**q**) and does not require running simulations of the perturbed ensemble. This allows us to use the fast PSO method ^51,52^ to find an optimal perturbation ***λ*** that minimizes the objective function (Step 2 in Fig. 2a). We then update the CG force field using Eq. 2 and simulate a new CG ensemble with the updated Gō network (Step 3 in Fig. 2a). Finally, as Eq. 5 provides only an approximate correction and the Gō model is coupled to the Martini force field, we iteratively repeat steps 2–3 until convergence (Step 4 in Fig. 2a).

To test the robustness of our method, we repeated the optimization three times for each test protein. In all cases, the optimizations converged within ∼30 steps, with the largest gains in agreement occurring within the first 15 steps (Fig. 2b and Fig. S3). This suggests that a typical optimization of the Gō network for a small to medium-sized protein only requires a few tens of microseconds of CG sampling to converge efficiently, and because our algorithm is highly parallelizable, the wall clock time can be reduced even further if more compute nodes are available.

We then tested how the choice of the initial (uniform) network strength affected the optimization results. We performed additional Gō network optimizations, starting from different *ε* values for the corresponding Lennard-Jones interactions: 10 kJ/mol, 12.5 kJ/mol, 15 kJ/mol, and 17.5 kJ/mol. Irrespective of the choice of the initial network strength or the test protein, we observed similarly fast and efficient convergence within ∼30 iterations. The resulting Gō networks yielded CG ensembles with SWD values near 0.1 nm relative to the reference all-atom MD ensembles (Fig. S3) – more than an order of magnitude improvement compared to the values reported for uniform Gō networks (Fig. 1d).

Optimization yielded unique Gō networks for each protein, with some bonds strengthening and others weakening (Fig. 2d). To assess these changes more systematically, we analyzed the distributions of the optimized interaction strengths (Fig. 2c). Optimization replicas started from the same initial network strengths converged to identical distributions (within the sampling error), whereas the distribution means differed for different starting strengths (Fig. 2c). These many minima and the flat optimization landscape are expected because the target observable, the essential dynamics, depends on collective motion patterns rather than on specific pairwise interactions. As a result, many different redistributions of interaction strengths can produce the same overall dynamical behavior. The stochastic optimization therefore explores a broad manifold of equivalent minima with comparable mean energies.

To compare the optimized ensembles more systematically, we computed (i) the Pearson correlation of the optimized bond energies and (ii) the *L*_2_-norm of the Laplacian eigenvalue spectrum across the different networks (Fig. 2e). The Pearson correlation and the *L*_2_-norm were 0.88 − 0.97 and 0.02 − 0.03, respectively, for the replicas started from the same initial network strength. When comparing replicas started from different initial strengths, the Pearson correlation ranged between 0.77 and 0.88 and the *L*_2_-norm between 0.04 and 0.08 (Fig. 2e and Fig. S6). These supports our hypothesis that the algorithm primarily reweights the relative energies of the Gō bonds to reproduce the essential dynamics, rather than converging toward a single optimal set of absolute bond strengths. This observation is additionally supported by analyzing the total potential energies of an optimized network. The total energy increases only for low initial network strengths (*e*.*g*., *ε* = 10 kJ/mol), whereas it remained largely unchanged for the optimizations started from high interaction strength (*e*.*g*., *ε* = 17.5 kJ/mol).(Fig. 2c and Fig. S4)

Although the optimization converges to multiple distinct parameter sets, these solutions are effectively equivalent with respect to the target observables. All reproduce the same essential dynamics and yield indistinguishable collective motions along the principal components. This degeneracy might formally resemble overfitting, but in this context it merely reflects the redundancy of the parameter space. The optimization is thus underdetermined rather than overfitted: different parameter sets encode the same large-scale motions equally well.

In short, the proposed optimization algorithm demonstrates fast convergence within a few tens of iterations, is robust with respect to the target of optimization and choice of the initial Gō network strength and the test protein, and produces CG ensembles that are in excellent agreement with the reference all-atom MD ensembles in terms of their essential dynamics.

### Optimized Gō networks reproduce essential atomistic dynamics and improve local protein flexibility

As stated in the first section, our aim is to improve the agreement between the reference atomistic and CG free energy landscapes in the essential subspace. To this end, we optimized the Gō networks for T4L, RBP, and MBP using the procedure described in the previous section (Fig. 2) and compared them to the corresponding unoptimized ensembles using a uniform Gō network (Fig. 3a). We set the interaction strength of the unoptimized networks to *ε* = 12.5 kJ/mol; however, a similar behavior was observed for the other tested interaction strengths (*ε* = 10.0−17.5 kJ/mol).

**Figure 3:**
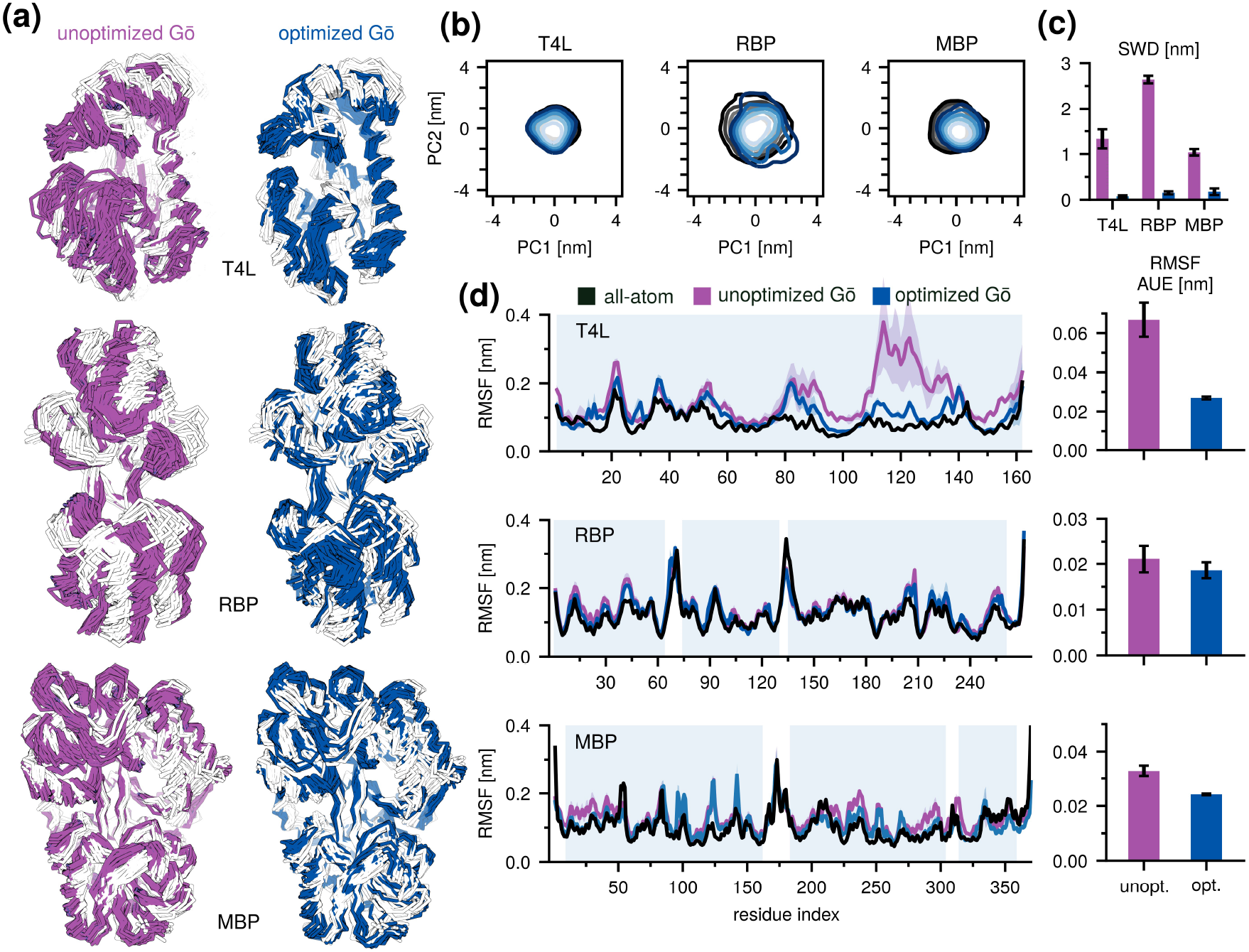
Performance of optimized GōMartini. **(a)** Representative ensembles for each protein generated with the unoptimized (left, purple) and optimized (right, blue) Gō network. Reference atomistic ensembles are depicted in white. **(b)** CG trajectories projected onto the all-atom essential subspace. Same color coding as in **(a) (c)** SWD values between the atomistic reference distribution and the unoptimized (purple) or optimized (blue) CG distributions projected onto the all-atom essential subspace.**(d)** Root mean squared fluctuation profiles for the unoptimized and optimized Gō networks compared to the reference atomistic profiles. Bar plots quantify the per-residue average unsigned error to the atomistic profiles.

Simulations with the unoptimized, uniform Gō network produced ensembles with excess fluctuations and partial unfolding (Fig. 3a). In contrast, a qualitative comparison of the optimized and unoptimized CG ensembles showed that the former were visually more similar to the reference all-atom ensembles – especially in terms of the alignment of small loops and the relative positions of motifs (Fig. 3a). A more detailed analysis of the essential subspaces revealed a significant improvement when comparing them to that of the reference all-atom ensembles. Specifically, we found that SWD decreased significantly from, on average, 1.67 ± 0.70 nm to 0.06 ± 0.02 nm for all tested protein systems (Fig. 3b,c and Fig. S7). RSMIP increased from 0.76 ± 0.16 to 0.83 ± 0.10, while CO improved from 0.58 ± 0.27 to 0.71 ± 0.17 (Fig. S8).

Having targeted the first three PCs in the optimization, we also calculated the RMSIP and CO over the first five and ten PCs. We found improvements relative to the unoptimized Gō network even when these additional PCs were included (Fig. S5).

In addition, we analyzed local root mean square fluctuations (RMSF) of the backbone beads of the optimized and unoptimized CG ensembles, as these are often a primary target in optimizing protein dynamics in Martini. Because it was not an explicit optimization target, the RMSF served as an excellent independent cross-validation metric, showing markedly improved agreement between the optimized CG and reference all-atom ensembles. Specifically, the per-residue average unsigned deviation decreased by roughly a half, from 0.04 ± 0.02 nm to 0.02 ± 0.00 nm (Fig. 3d), suggesting that our global optimization approach can simultaneously improve the local fluctuation profile.

To test whether improving local RMSF profiles alone enhances agreement in the essential subspace, we analyzed unoptimized ensembles with uniformly increased interaction strengths. Although higher uniform interaction strength reduced bead fluctuations and improved RMSF, this stiffening led to only minor gains in overlap with the atomistic essential subspace (Fig. S8). In contrast, our optimization achieved markedly better agreement in both the fluctuation profile and essential subspace with much smaller perturbations to the total interaction energy, indicating that reproducing local RMSF does not necessarily ensure accurate global dynamics.

## Conclusions

Here, we developed an automated optimization framework, *PoGō*, that combines a low-dimensional representation of protein dynamics, an efficient global optimization algorithm (PSO), and analytical perturbation theory to fine-tune the interaction strengths of Gō -based networks. Validation on three proteins with varying complexity and essential dynamics demonstrated that the method achieves excellent agreement with atomistic reference ensembles within only few iterations.

In contrast, the standard Martini force field—supported by either the default elastic network or a uniform-strength Gō network—was unable to accurately reproduce atomistic essential dynamics. Increasing the network interaction strengths or deriving equilibrium bead–bead distances from all-atom simulations yielded only marginal improvements, with no universal parameter set performing consistently across systems.

Despite extensive work highlighting the importance of protein essential dynamics, ^12,14,20–24,26,28^ these features are often overlooked when optimizing CG models, where more emphasis is typically placed on reproducing local fluctuations. ^43^ While improving essential dynamics can naturally enhance local fluctuations, the reverse is not necessarily true. Our results demonstrate that targeting the lowest-energy collective motions might provide a more efficient route to achieving both accurate global and local dynamics.

Our method is not restricted to a particular reaction coordinate. Although in this study we focused free-energy landscapes within the essential subspace, the same framework could be used to optimize probability distributions along any generalized reaction coordinate, such as the radius of gyration or the distance between the centers of mass of two protein domains. The dimensionality of the essential subspace is also tunable; however, using more than five dimensions bears the risk of undersampling in all-atom molecular dynamics (MD) simulations and makes optimization challenging due to rapidly diminishing probabilities in high-dimensional space. Furthermore, the approach is not limited to the Lennard–Jones potentials of GōMartini and is compatible with any additive CG force field, rendering it a general and transferable strategy for global ensemble refinement in high-dimensional parameter spaces.

Despite its broad applicability and flexibility, several limitations should be considered. First, the current implementation optimizes only the depths of the Lennard–Jones potentials in the Gō network, requiring any additional parameters (e.g., equilibrium bead–bead distances) to be known and fixed prior to optimization. Second, the network topology remains fixed throughout the procedure. Selecting an optimal topology is a complex problem in its own right and lies beyond the present scope. Nevertheless, too dense network or high initial network strength may hinder studies involving mutations or ligand binding where local flexibility is essential. Alternative network generation methods such as OLIVES ^63^ may provide more suitable starting points in such cases. Third, the optimization quality depends on the quality of the target ensemble. Although we employ all-atom MD references here, the target ensemble might also be derived from deep learning approaches (e.g., BioEmu ^64^) or from experimental data. ^65^

In conclusion, we describe a generalizable, automated framework for optimizing CG models to reproduce atomistic protein dynamics. Integrating physics-inspired ^63^ or machine-learned priors ^66,67^ for network topology design could further enhance performance. At the same time, incorporating experimentally derived ensemble information—from techniques such as electron microscopy, ^65,68^ SAXS, or NMR—offers a promising avenue for integrative structural biology. Altogether, this framework enables more reliable and high-throughput studies of protein dynamics, functional mechanisms, mutational effects, and protein assembly within CG representations.

## Contributions

M.K. conceptualized the project and implemented the initial optimization approach. M.K. and C.J.W. developed the final optimization protocol, performed the simulations, analyzed the data, and drafted the manuscript. H.G. and M.I. secured funding and supervised the project. All authors reviewed and edited the final manuscript.

## Acknowledgements

All authors acknowledge the support provided by the Max Planck Society. M.K., M.I., and H.G. additionally thank the Deutsche Forschungsgemeinschaft (DFG): Project-ID 449750155 – RTG 2756, Project A1 for supporting this research. C.J.W. is grateful to the National Sciences and Engineering Research Council of Canada (NSERC) (PGSD-599306-205) for funding. Open access funding was provided by the Max Planck Society.

## Code availability

The *PoGō* optimization code is available at https://github.com/wilsonjcarter/pogo.

## Notes

### Competing Interest Statement

The authors have declared no competing interest.

