## Supporting Information for "Improving conformational ensembles of folded proteins in GōMartini"

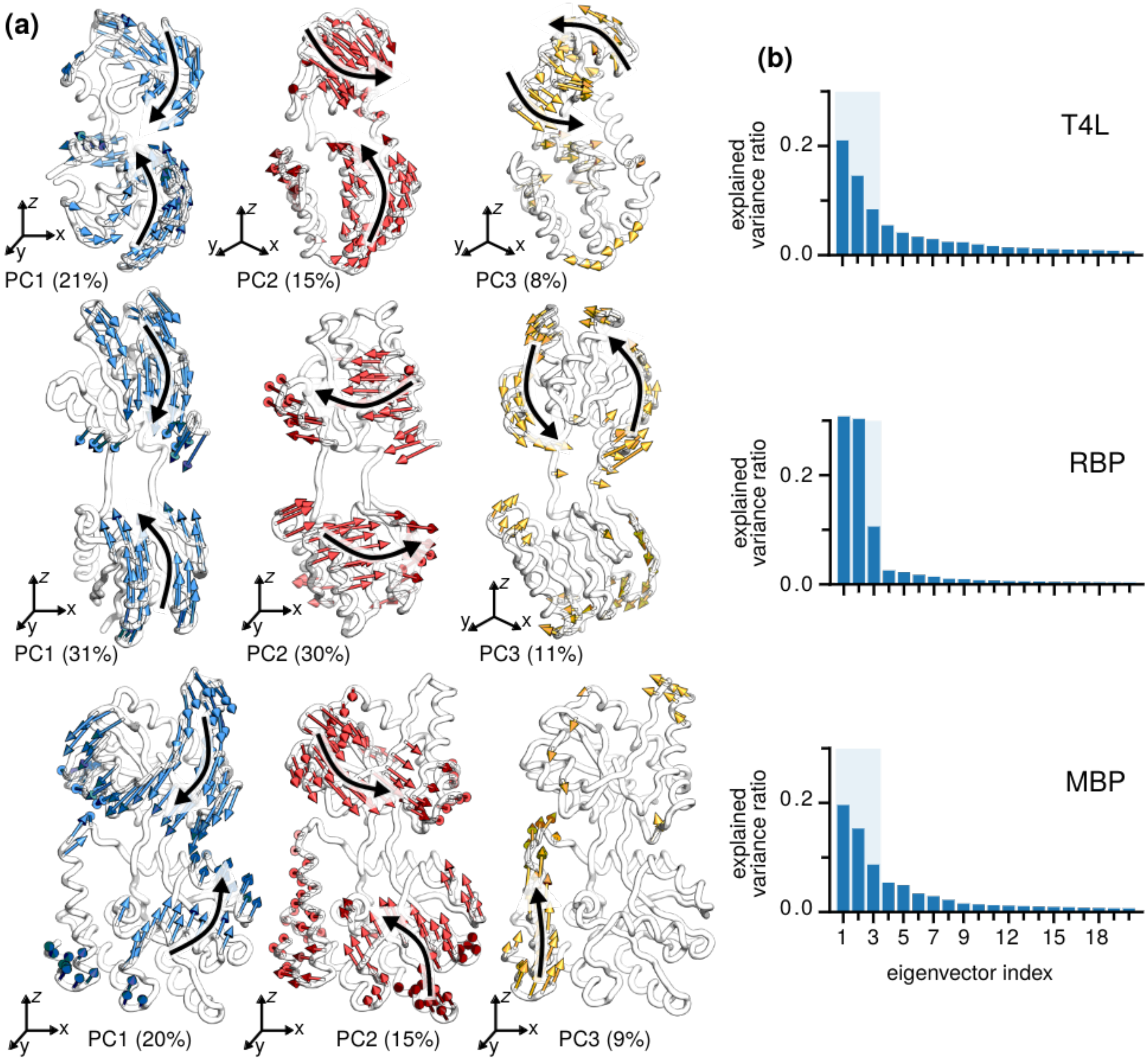


Figure S1: **(a)** Projected motions of the first three principal components. **(b)** Explained variance of the first twenty components.

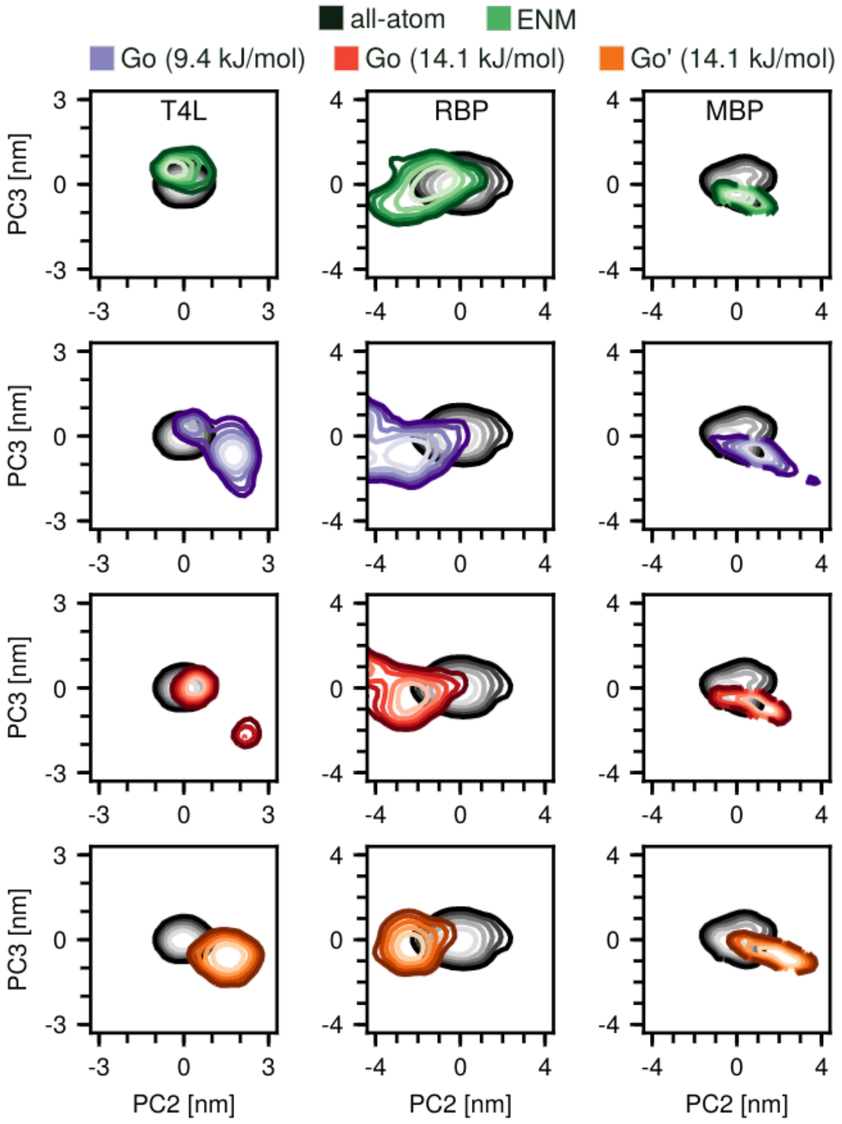


Figure S2: All-atom MD and CG trajectories projected onto the essential subspace of the MD ensembles (PC2 and PC3). Each column corresponds to a different protein, and each row corresponds to a different CG setup.

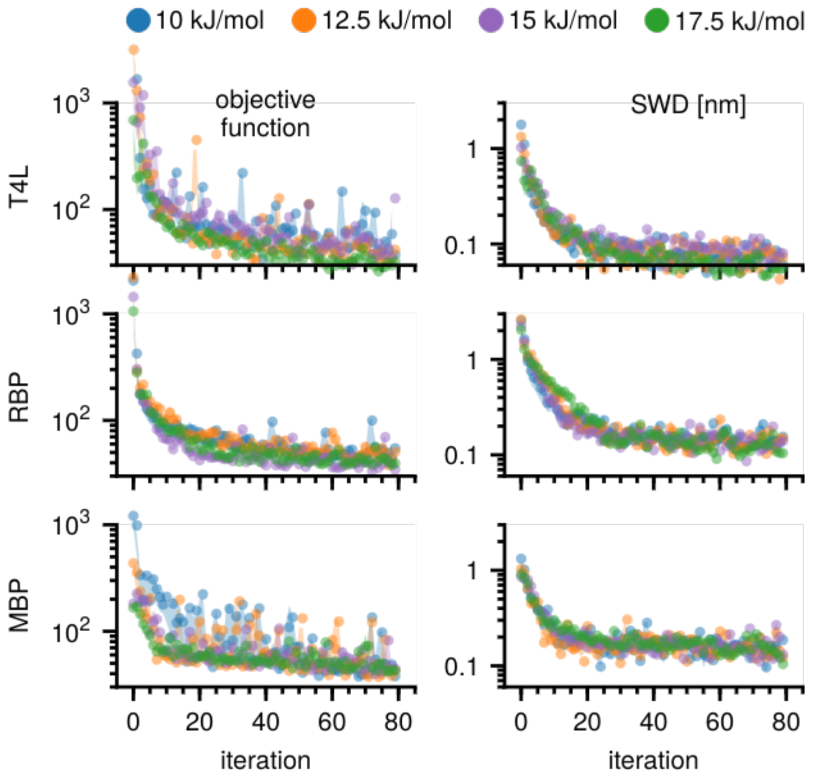


Figure S3: Convergence behavior for optimizations started with different uniform network strengths.

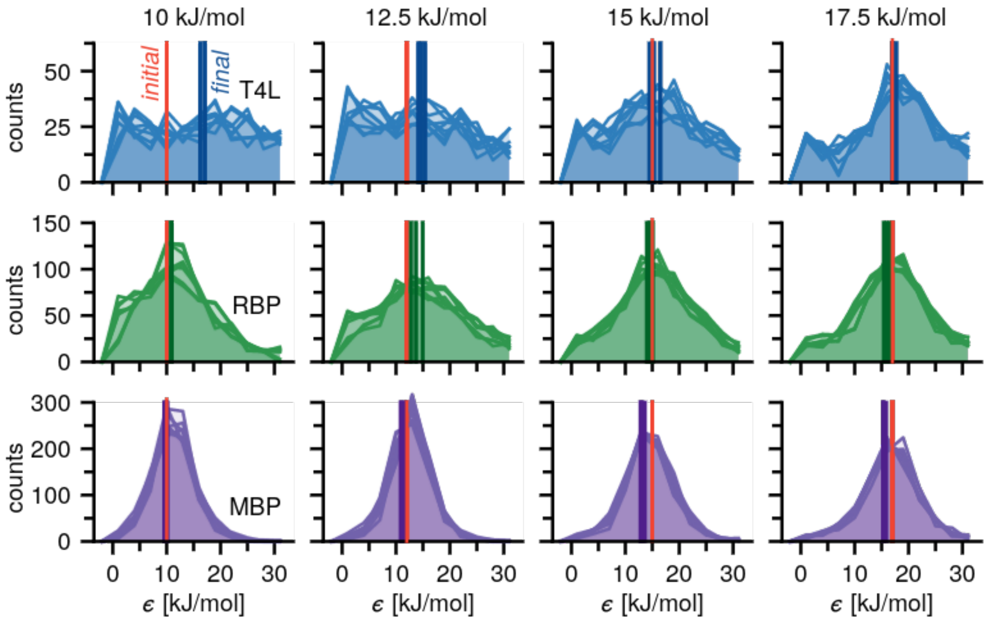


Figure S4: Distribution of the optimized Gō network edge weights as a function of the starting network strength. Initial and final averages are indicated with vertical lines.

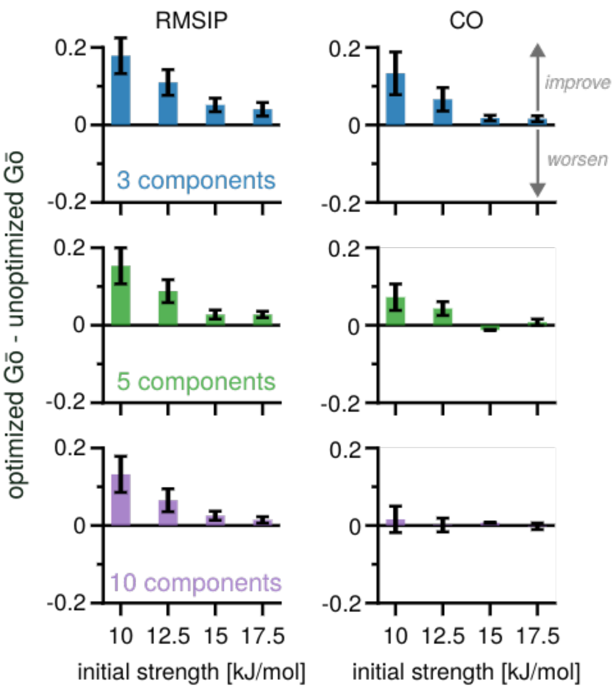


Figure S5: Change in the RMSIP and CO between the unoptimized and optimized Gō networks averaged over proteins and replicates. 3, 5, and 10 principal components were considered for the calculations. Positive values indicate improved agreement with the atomistic reference over unoptimized while negative values indicate worse agreement.

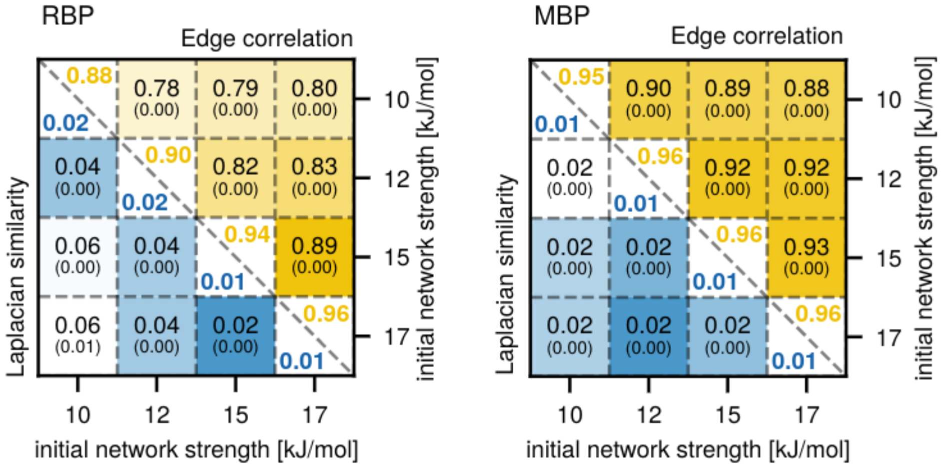


Figure S6: Correlation properties of Gō networks initialized from different uniform strengths. Standard errors are indicated in brackets. Numbers on the diagonal correspond to intra-replica metrics.

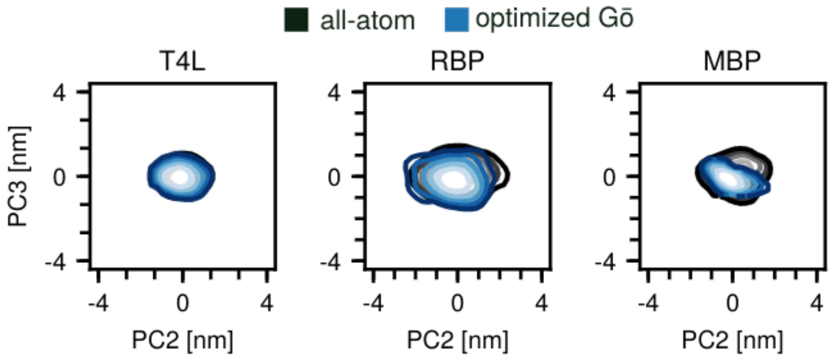


Figure S7: All-atom MD and optimized CG trajectories projected onto the essential subspace of the MD ensembles (PC2 and PC3). Each column corresponds to a different protein.

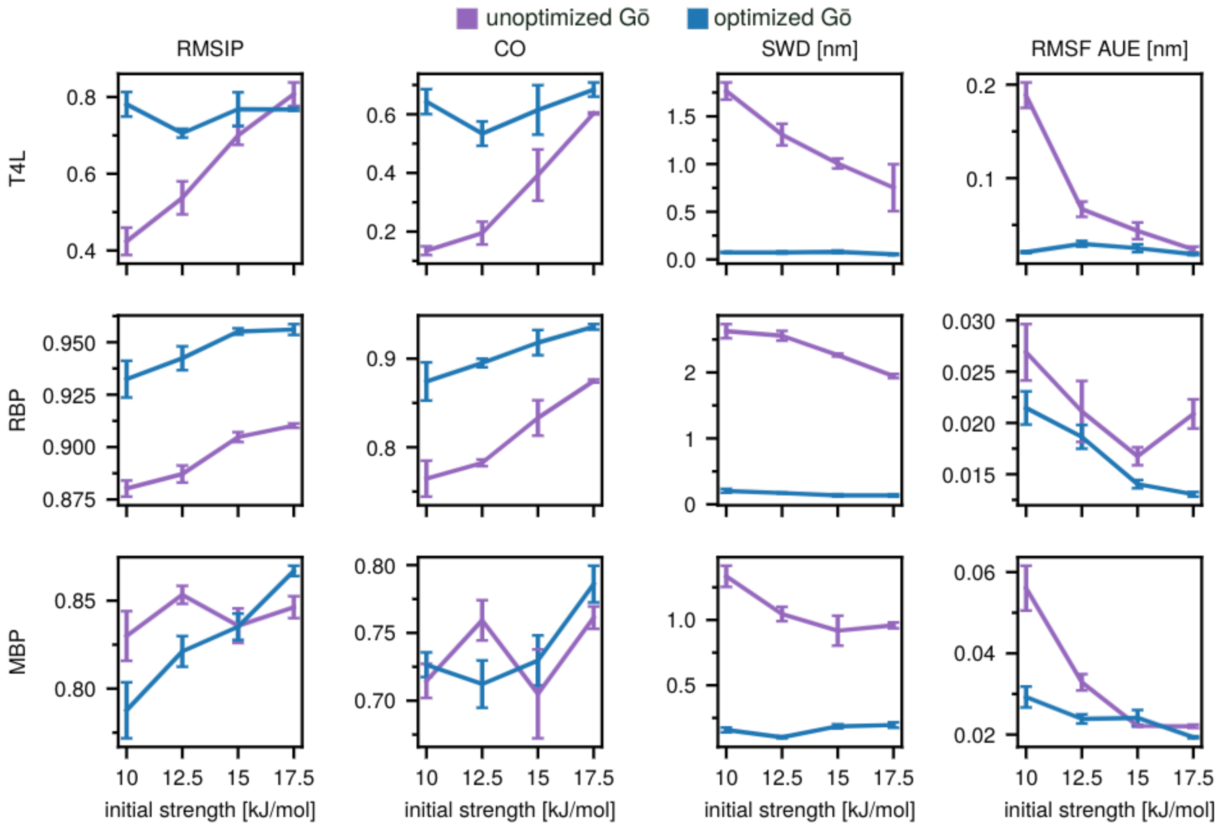


Figure S8: Impact of the initial Gō network strength on both unoptimized (purple) and optimized (blue) structural metrics for T4L (upper row), RBP (middle row) and MBP (bottom row). Error bars correspond to three independent repeats.
